# Fine-grained mapping of cortical somatotopies in chronic Complex Regional Pain Syndrome

**DOI:** 10.1101/409094

**Authors:** Flavia Mancini, Audrey P Wang, Mark M. Schira, Zoey J. Isherwood, James H. McAuley, Giandomenico D Iannetti, Martin I. Sereno, G. Lorimer Moseley, Caroline D. Rae

**Author notes:** Shared contribution (Flavia Mancini, Audrey P Wang). Corresponding author: Flavia Mancini University of Cambridge, Department of Engineering, Computational and Biological Learning, Trumpington Street, Cambridge, CB2 1PZ.

## Abstract

It has long been thought that severe chronic pain conditions, such as Complex Regional Pain Syndrome (CRPS), are not only associated with, but even maintained by a reorganisation of the somatotopic representation of the affected limb in primary somatosensory cortex (S1). This notion has driven treatments that aim to restore S1 representations, such as sensory discrimination training and mirror therapy. However, this notion is based on both indirect and incomplete evidence obtained with imaging methods with low spatial resolution. Here, we used functional MRI to characterize the S1 representation of the affected and unaffected hand in patients with unilateral CRPS. At the group level, the cortical area, location, and geometry of the S1 representation of the CRPS hand were largely comparable to those of the healthy hand and controls. However, the area of the map of the affected hand was modulated by disease duration (the smaller the map, the more chronic the CRPS), but not by pain intensity, pain sensitivity and severity of the physical disability. Thus, if any map reorganization occurs, it does not appear to be directly related to our pain measures. These findings compel us to reconsider the cortical mechanisms underlying CRPS and the rationale for interventions that aim to “restore” somatotopic representations to treat pain.

**Significance statement:** This study shows that the spatial map of the fingers in S1 is largely preserved in chronic CRPS. Shrinkage of the area of the affected hand map can occur in the most chronic stages of disease. Map shrinkage is related to CRPS duration rather than diagnosis, and is unrelated to how much pain patients experience or to the severity of the physical disability. These findings challenge the rationale for using sensory interventions to treat pain by restoring somatotopic representations in CRPS patients.

## Introduction

Chronic pain is a highly common and debilitating disorder, that can be associated with functional and morphological changes in the brain. For instance, it has long been thought that some severe chronic pain conditions, such as Complex Regional Pain Syndrome (CRPS), are not only associated with, but even maintained by, maladaptive topographic changes in the primary somatosensory cortex (S1) (Maihofner et al., 2003, 2004). Magneto- and electro-encephalography (MEG, EEG) studies have suggested that the representation of the CRPS hand in S1 is abnormally smaller than the cortical representation of the healthy hand (Juottonen et al., 2002; Maihofner et al., 2003; Pleger et al., 2004; Vartiainen et al., 2008; Vartiainen et al., 2009). The notion of S1 reorganisation has been central to our understanding of the condition (Marinus et al., 2011) and has driven physiotherapy interventions aimed at restoring sensorimotor representations of CRPS limbs, such as mirror-visual feedback (McCabe et al., 2003; Smart et al., 2016) and sensory discrimination training (Pleger et al., 2005; Moseley et al., 2008a). Here, we revisit the notion of S1 reorganisation with the better tools that modern functional MRI currently offers: high spatial resolution and phase encoded methods that provide reliable and unbiased measures of the cortical somatotopy of the hand (Mancini et al., 2012; Sanchez-Panchuelo et al., 2012; Kolasinski et al., 2016a).

In all previous studies on CRPS, the size of the hand map was estimated both *indirectly and incompletely*: it was estimated by measuring the Euclidean distance between activation loci of the thumb or index finger relative to that of the little finger. The somatotopy of the full hand has never been characterized in CRPS patients. A more reliable fMRI method for studying cortical topographic representations is based on phase-encoded mapping, which reveals the spatial preference of cortical neural populations (Sereno et al., 1995; Silver and Kastner, 2009; Sereno and Huang, 2014). This method involves delivering a periodic sensory stimulus to different portions of the receptive surface and evaluating which voxels selectively respond to the spatial frequency of the stimulation. Voxels sensitive to the stimulus respond when the stimulus passes through the preferred spatial location and decay as the stimulus moves away (Chen et al., 2017). The response phase angle, extracted using a Fourier transform (Mancini et al., 2012), indicates the location preference for each voxel—in other words, the position of the receptive fields of the population of neurons sampled by the voxel.

Using phase-encoded mapping, we provide the first complete characterisation and quantification of the representation of the fingers (i.e. with exclusion of the thumb) in patients with chronic and unilateral CRPS to the upper limb. We tested whether the S1 representation of the fingertips of the affected hand was different from that of the healthy hand of CRPS patients and from controls in terms of its spatial extent, location relative to the central sulcus, and geometry (i.e. variability of the map gradients).

## Materials and Methods

### Participants

We recruited 20 adults with unilateral CRPS to the upper limb and 20 healthy controls (HC) matched for age, gender and handedness. Each participant gave written informed consent to take part in the study. All experimental procedures were carried out in accordance with the Declaration of Helsinki and approved by both the Human Research Ethics Committee of the University of New South Wales (HC13214) and by the Human Ethics Committee of the South Eastern Local Health District (HREC 10/051). Inclusion criteria for control participants were: (1) pain-free at that time of the study; (2) no prior history of a significant chronic pain, psychiatric or medical disorder; (3) no history of substance abuse. Inclusion criteria for CRPS patients were: (1) a diagnosis of unilateral CRPS to the upper limb or hand according to the Budapest research criteria (Harden et al., 2010); (2) CRPS duration greater than 3 months; (3) no history of substance abuse and no psychiatric-comorbidities. Five of 40 participants were excluded from the study due to the following problems: MRI scanner failure (subject #31) or acquisition problems (#13, #15, #32) and a control participant reported pain to the wrist on the day of scan (a median nerve compression was subsequently diagnosed, #42). The demographic and clinical information of the remaining sample (Controls: n = 17; CRPS to the left hand: n = 8; CRPS to the right hand: n = 10) is reported in Table 1.

**Table 1.** *Demographic and clinical information of the study sample.* Range of motion, motor weakness, tremor, allodynia: ‘-‘ indicates no abnormality, whereas ‘+’ indicates presence of a symptom. Intensity of pain to the upper limb during scans were evaluated on a Likert scale from 0 (no pain) to 10 (worst pain imaginable). PPT indicates the Pressure Pain threshold and is reported in kg/cm^2^ units. The laterality score is derived from the Edinburgh Handedness Inventory and ranges from -100 (left-hand dominant) to +100 (right-hand dominant).

| ID | Group | Age | Gender | CRPS Duration (y) | Incident at onset | Range of Motion | Motor weakness | Tremor | Allodynia | Pain rating during scan | PPT left hand | PPT right hand | Laterality score |
| --- | --- | --- | --- | --- | --- | --- | --- | --- | --- | --- | --- | --- | --- |
| 4 | Control | 38.5 | F | n/a | n/a | - | - | - | - | 1 | 50.5 | 62.2 | 87.5 |
| 5 | Control | 42.8 | M | n/a | n/a | - | - | - | - | 4 | 13.4 | 13.6 | 87.5 |
| 7 | Control | 52.8 | M | n/a | n/a | - | - | - | - | 0 | 43.7 | 40.8 | 87.5 |
| 10 | Control | 41.1 | M | n/a | n/a | - | - | - | - | 0 | 4.72 | 3.87 | 100 |
| 11 | Control | 56.6 | M | n/a | n/a | - | - | - | - | 0 | 5.03 | 4.83 | 73.3 |
| 16 | Control | 42.3 | M | n/a | n/a | - | - | - | - | 0 | 3.6 | 4.78 | 87.5 |
| 21 | Control | 34.1 | M | n/a | n/a | - | - | - | - | 0 | 3.84 | 4.72 | 64.7 |
| 22 | Control | 53.0 | M | n/a | n/a | - | - | + | - | 0 | 5.36 | 5.55 | 66.7 |
| 27 | Control | 48.7 | F | n/a | n/a | - | - | - | - | 0 | 7.92 | 7.39 | 100 |
| 30 | Control | 56.5 | M | n/a | n/a | - | - | - | - | 0 | 3.92 | 3.27 | 83.3 |
| 34 | Control | 46.9 | M | n/a | n/a | - | - | - | - | 0 | 5.58 | 5.73 | 100 |
| 35 | Control | 38.4 | M | n/a | n/a | - | - | - | - | 0 | 3.7 | 4.75 | 100 |
| 36 | Control | 47.8 | M | n/a | n/a | - | - | - | - | 0 | 5.6 | 5.53 | -100 |
| 37 | Control | 25.2 | F | n/a | n/a | - | - | - | - | 0 | 4.41 | 4.88 | 100 |
| 38 | Control | 19.9 | M | n/a | n/a | - | - | + | - | 0 | 5.4 | 5.14 | 12.5 |
| 39 | Control | 49.2 | M | n/a | n/a | - | - | - | - | 0 | 6.52 | 7.71 | 100 |
| 40 | Control | 69.4 | M | n/a | n/a | - | - | - | - | 0 | 3.83 | 3.42 | 100 |
| 6 | CRPS to left hand | 42.4 | M | 1.2 | Wrist fracture | + | + | + | + | 7 | 1.9 | 12.2 | 50 |
| 9 | CRPS to left hand | 42.8 | M | 0.9 | Wrist injury | + | + | - | + | 8 | 10.6 | 12.3 | 83.3 |
| 12 | CRPS to left hand | 55.6 | M | 0.5 | Frozen shoulder | + | + | + | - | 9 | 2.06 | 4.13 | 83.3 |
| 14 | CRPS to left hand | 45.3 | M | 3.8 | Hand surgery | + | + | + | + | 7 | 1.8 | 2.55 | 4.3 |
| 23 | CRPS to left hand | 66.7 | M | 0.4 | Hand injury and infection | + | + | + | + | 2 | 0.88 | 4.31 | 73.9 |
| 26 | CRPS to left hand | 47.8 | M | 4.1 | Hand surgery | - | + | + | + | 0 | 2.9 | 3.89 | 100 |
| 28 | CRPS to left hand | 41.9 | M | 14.9 | Hand trauma injury | + | + | + | + | 7 | 0.33 | 3.55 | 100 |
| 29 | CRPS to left hand | 29.2 | M | 1.8 | Hand injury | + | + | + | + | 6 | 0.76 | 2.06 | -100 |
| 3 | CRPS to right hand | 53.4 | M | 4.5 | Shoulder injury | - | + | - | + | 9 | 49.2 | 15.5 | 100 |
| 8 | CRPS to right hand | 38.1 | M | 0.4 | Hand and wrist injury | + | + | + | + | 8 | 6.925 | 2.925 | 85.7 |
| 17 | CRPS to right hand | 51.6 | M | 0.4 | Hand injury | + | + | + | + | 2 | 4.77 | 3.53 | 89.5 |
| 18 | CRPS to right hand | 34.7 | M | 2.9 | Hand fracture | + | + | + | + | 7 | 3.82 | 1.3 | 73.3 |
| 19 | CRPS to right hand | 48.7 | F | 2.6 | Wrist fracture and surgery | + | + | + | + | 8 | 12.59 | 4.94 | 100 |
| 20 | CRPS to right hand | 46.7 | M | 7.5 | Shoulder injury | - | + | + | + | 5 | 2.37 | 1.06 | 66.7 |
| 24 | CRPS to right hand | 56.8 | M | 14.6 | Arm injury | + | + | + | + | 10 | 3.07 | 0.9 | 100 |
| **25** | CRPS to right hand | 26.6 | M | 1.9 | Wrist fracture | + | + | + | + | 7 | 3.1 | 0.98 | -66.7 |
| **33** | CRPS to right hand | 21.4 | F | 2.6 | Wrist and Hand trauma injury with surgery | + | + | + | + | 0 | 4.79 | 3.84 | 100 |
| **41** | CRPS to right hand | 44.9 | M | 18.3 | Road traffic accident | + | + | + | + | 9 | 3.63 | 2.45 | 64.7 |

### Clinical evaluation

Patients were clinically evaluated according to the Budapest research criteria (Harden et al., 2010) by a blinded assessor of the research team on the first session of the study to confirm that the research criteria were met. As part of the clinical and diagnostic assessment of CRPS, we assessed pressure pain thresholds (PPT; kg/cm^2^) using a digital pressure algometer (Wagner instrument, Greenwich, USA) on two sites of each hand: the thenar eminence and the third proximal interphalangeal joint. Pain intensity was also rated using an 11-points Likert scale, where 0 corresponded to “no pain” and 10 indicated “the worst pain imaginable, like a red hot poker through your eye”. The intensity of spontaneous pain in the upper limb was rated in all patients immediately before, during and after the imaging session. Two control participants reported discomfort and mild to moderate postural pain to the upper limb during the scanning session (Table 1). Furthermore, the *Quick*Dash (Disabilities of the Arm, Shoulder and Hand) questionnaire was administered to all participants; the *Quick*Dash measures physical function and symptoms in people with musculoskeletal disorders of the upper limb (Kennedy et al., 2011).

### Stimuli

We used a customised stimulus (polypropylene probe with a rounded tip) because CRPS skin physiology and symptoms (hand dystonia, pain) preclude the use of conventional and automated mechanical stimulation for the prolonged time required for phase-encoded mapping of the fingertips (approx. 40 minutes). For example, hand dystonia makes it difficult to target the same skin regions with air-puffs throughout the imaging session; this would have resulted in scan quality deterioration or early scan termination. CRPS-related hyperhidrosis (i.e. excessive sweating) precludes the use of electrical and vibrotactile stimulation for a long time.

All control participants reported the stimulus as being clearly detectable, neither painful nor unpleasant, and similarly intense on the different fingers of the two hands. All patients described the sensation that was elicited by stimulation of the unaffected fingers, in similar terms to those used by the control participants. Patients described the sensation that was elicited by stimulation of the affected fingers in a variety of ways; “burning”, “tingling”, “pain”, “brushing like with a sharp object”, “horrible”, “itchy”, “scraping”, “like a needle prick”, “electric shooting pain”. These terms are consistent with the clinical phenomenon of allodynia.

Participants did not report systematic differences in stimulus perception across the fingertips of the same hand. Pain intensity fluctuates over time in most chronic pain conditions (including CRPS), even despite highly-controlled and reproducible stimulation (Foss et al., 2006). However, such fluctuations are unlikely to confound our measures of cortical somatotopy. Indeed, our analysis method allowed to dissect the magnitude of the brain responses from their spatial organization. All our analyses did not focus on the magnitude of the S1 responses, but on their spatial organisation, which is not confounded by unavoidable fluctuations of perceived stimulus intensity in CRPS patients.

### Procedure

Each participant laid supine inside the scanner bore with both hands palm upwards. Participant’s arms and hands were propped with cushions and pads to minimise movements. The stimulus consisted of periodic stimulation of the fingertips of both hands. In each stimulation cycle, the tips of the index, middle, ring, and little fingers were successively stimulated using a customised probe (see below). Each fingertip was stimulated for 6 s, and each cycle (four fingers x 6 s = 24 s) was interleaved by 6 s of rest. Twelve cycles were administered in each of the four consecutive functional runs (approx. 10 minutes each). Two trained experimenters stimulated the tips of homologous fingers of the right and left hands simultaneously. The experimenters received auditory cues through headphones, synchronising the location and timing of each stimulus. The thumb was not stimulated to reduce scanning time and due to practical difficulties in stimulating the thumb in succession to the other fingertips (patients could not keep the hand open flat for prolonged periods of time).

Our choice of bilateral stimulation was motivated by the need to map the fingertips of both hands in a single imaging session (several patients travelled from distant regions in Australia). Importantly, our choice was grounded on neuroscientific evidence that there are extremely limited trans-callosal connections between the hand representations of S1 in the primate brain (Jones and Hendry, 1980; Killackey et al., 1983). We note that some studies have reported an inhibitory response to hand stimulation in ipsilateral S1 (Hlushchuk and Hari, 2006; Lipton et al., 2006; Klingner et al., 2011). The deactivation of ipsilateral S1 is most likely mediated by an input that ascends the contralateral pathway to a higher-order cortical area, crosses in the corpus callosum, and is then fed back to area 3b in S1 (Lipton et al., 2006; Tommerdahl et al., 2006). Ipsilateral finger representations are engaged in active movement, but not during somatosensory processing (Berlot et al., 2018). Peripheral nerve injury can enhance activity in ipsilateral S1 (Fornander et al., 2016), especially at the level of interneurons in laminae V and VI (Pelled et al., 2009). Crucially, ipsilateral activations and deactivations in S1 are diffused and not somatotopically specific (Helmich et al., 2005; Reed et al., 2011; Ann Stringer et al., 2014; Geva et al., 2017). They can modulate the amplitude of the S1 response (which is not of interest here), but there is no evidence that they affect the spatial (somatotopic) organisation of the contralateral responses (Reed et al., 2011; Ann Stringer et al., 2014; Geva et al., 2017). This is further confirmed by our preliminary imaging data, in which we found that unilateral vs. bilateral fingertip mapping yielded both greatly similar and highly reproducible fingertip maps in S1 (Figure 1). Therefore, we considered bilateral finger stimulation as a resource-efficient method to map the S1 somatotopy of the fingers of both hands in a single imaging session, thus boosting recruitment and compliance of CRPS patients.

**Figure 1.**
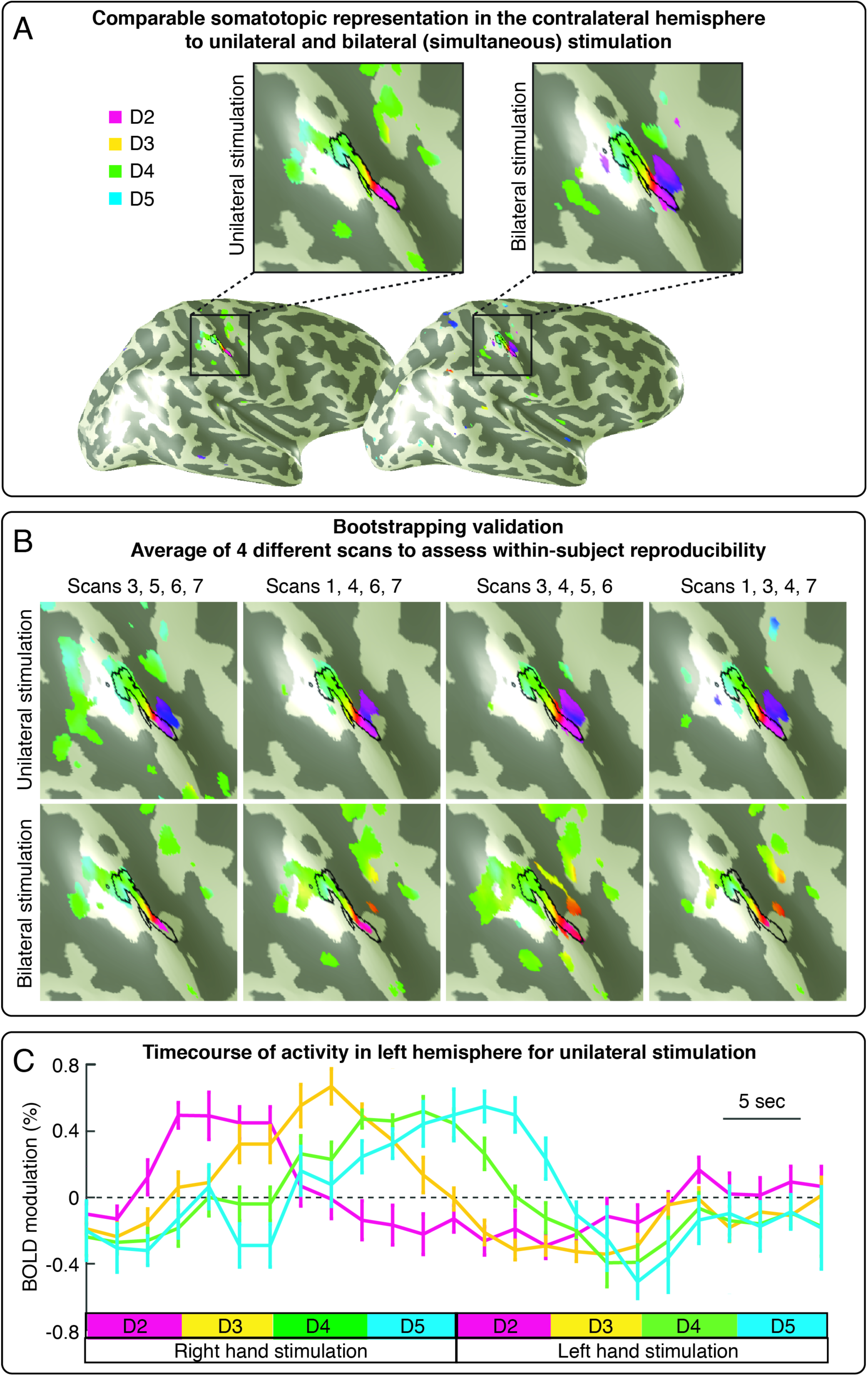
*Preliminary results that guided the design of the finger mapping protocol. (A) Comparable somatotopic representation in the contralateral S1 to unilateral and bilateral finger stimulation, at within-subject level.* The map of the fingertips (d2-d5) in contralateral S1 was strikingly similar in a condition in which we stimulated the fingertips of one hand at time and in another condition whereby we stroked homologous fingertips of both hands simultaneously. *(B) Bootstrapping validation.* We validated the results shown in panel A using a bootstrapping approach. Seven functional runs per condition (unilateral stimulation, bilateral stimulation) were collected in a single participant, in multiple scanning sessions. We selected, both recursively and randomly, 4 runs among the 7 collected per condition and averaged results across these 4 runs to assess intra-individual map reproducibility. The maps of the fingertips were highly reproducible in both unilateral and bilateral stimulation conditions. *(C) Time course of activity in the left hemisphere during unilateral fingertip stimulation.* Panel C shows the percent modulation of BOLD response in the left S1 induced by periodic stimulation of the fingertips of the right hand and left hand. We did not observed a spatially-tuned activation of the left S1 induced by left-hand stimulation.

### MRI acquisition

Echoplanar images (1.5 mm^3^ isotropic resolution, 183 volumes/run, 32 axial slices, flip angle = 82°, TR = 2s) were collected in four runs on a Philips Achieva TX 3T MRI scanner using a 32-channel head coil. FreeSurfer (https://surfer.nmr.mgh.harvard.edu/) was used to reconstruct the cortical surface for each subject from a structural T1 image (0.727×0.727 mm^2^ in-plane, 0.75 mm thick slices, 250 slices, flip angle = 8°, TR = 6.318 ms). In four subjects (28, 29, 33, 34), structural T1 images were corrected for non-uniform intensity using the AFNI’s tool ‘3dUnifize’ (https://afni.nimh.nih.gov), before surface reconstruction, because these images contained shading artefacts that could have affected segmentation.

### First-level MRI analyses

All first-level analyses were performed by a researcher (FM) blinded to the group condition (right CRPS, left CRPS, control). The first 3 volumes from EPIs were discarded from all analyses. Functional series were aligned and motion-corrected using the AFNI program ‘3dvolreg’. Using this as a starting point, functional-to-high resolution alignment was then refined using manual blink comparison using an adaptation of Freesurfer’s TkRegister implemented in csurf (http://www.cogsci.ucsd.edu/~sereno/.tmp/dist/csurf). After linear trend removal, aligned data from the four runs were raw-averaged, and then analysed using a fast Fourier transform, computed for the time series at each voxel fraction (vertex): this resulted in complex-valued signals with the phase angle and magnitude of the BOLD response at each voxel. The phase angle is the measure of interest here, because it reflects the spatial preference of a given voxel. Both Fourier and statistical analysis were performed using csurf. No spatial smoothing was performed before statistical analyses. Very low temporal frequencies and harmonics (< 0.005 Hz) were excluded because movement artefacts dominate responses at these frequencies, a procedure virtually identical to regressing out signals correlated with low frequency movements. High frequencies up to the Nyquist limit were allowed (i.e. half the sampling rate); this corresponds to no use of low pass filter. For display, a vector was generated whose amplitude is the square root of the F-ratio calculated by comparing the signal amplitude at the stimulus frequency to the signal at other noise frequencies and whose angle was the stimulus phase. To minimize the effect of superficial veins on BOLD signal change, superficial points along the surface normal to each vertex were disregarded (top 20% of the cortical thickness).

The F-ratio was subsequently corrected at p < 0.01 using a surface-based cluster correction for multiple comparisons as implemented by surfclust and randsurfclust within the csurf FreeSurfer framework (Hagler et al., 2006). The Fourier-transformed data were then sampled onto the individual cortical surface. Using this statistical threshold, we cut a label containing all vertices that showed a significant periodic response to finger stimulation (see one example in Figure 7A) and was localised within S1 (i.e. within the boundaries of areas 3a, 3b and 1, as estimated by the cortical parcellation tools implemented in Freesurfer). This label, or region of interest (ROI), is used as the input for the analyses described in the next sections. The phase-encoded stimulation procedure that we used is designed to map the hand region across fingers, not within fingers (Sanchez-Panchuelo et al., 2012). Therefore, we could not derive accurate ROIs for each finger in isolation. This is because voxels that are activated by more than one finger are masked out. Furthermore, we did not derive ROIs for the different subdivisions of S1 because a precise and reliable parcellation of the cortical surface at single-subject level would require microstructural imaging.

In a few cases, we could not identify any ROI with a response to fingertip stimulation (no response to either fingertip stimulation), even at uncorrected p < 0.05: subject #3, right hemisphere (patient with right CRPS); subject #20, left hemisphere (right CRPS); subject #24, left hemisphere (right CRPS); subject #28, left hemisphere (left CRPS); subject #29, right hemisphere (left CRPS). These cases were excluded from further analysis.

### Evaluation of hand map area

We calculated the surface area of the left- and right-hand maps, from each participant ROI. This was done after resampling the phase maps onto the original average brain volume, to control for inter-individual variability in brain size. In order to increase statistical power, we pooled data from the two CRPS groups and compared map area in the affected vs unaffected sides with both a frequentist and a Bayesian mixed-effects ANOVAs with a within-subject factor ‘side’ (2-levels: affected, unaffected) and a between-subjects factor ‘group’ (2-levels: controls, CRPS). In the CRPS group, we tested whether the area of the maps of the affected and unaffected hands could be explained using Bayesian linear regression models by the following variables: (1) CRPS duration; (2) the severity of upper limb disability as measured by the *Quick*Dash score; (3) pain intensity rating collected during the imaging session; (4) a severity score derived from the difference of PPT thresholds in the two hands as follows:

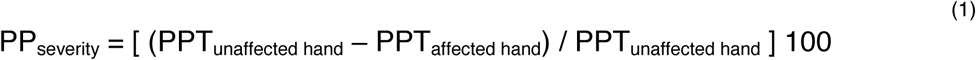

### Evaluation of hand map location

We controlled for individual differences in brain morphology as follows. We first inflated each participant’s cortical surface to a sphere and then non-linearly morphed it into alignment with an average spherical cortical surface using FreeSurfer’s tool mri_surf2surf (Fischl et al., 1999). This procedure maximizes alignment between sulci (including the central sulcus), while minimizing metric distortions across the surface. We resampled phase maps onto this average spherical surface (Freesurfer’s fsaverage) and calculated the location of the centroid of the map on this average surface. We investigated whether the map centroid was different across sides and groups, in two ways.

First, we tested whether the distribution of spherical coordinates was different across conditions (‘side’ and ‘group’). As a basis for this comparison, we used the Fisher probability density function (Fisher, 1953), which is the spherical coordinate system analogue of the Gaussian probability density function. This approach has been commonly used in the field of paleomagnetism and has also been applied for the analysis of direction data from diffusion tensor imaging (Hutchinson et al., 2012). We calculated the F statistics for the null hypothesis that sample observations from two groups are taken from the same population. The following equation was derived from Watson (Watson, 1956; Hutchinson et al., 2012) and used to compare two groups with N_1_ and N_2_ observed unit vectors and resultant vectors of length R_1_ and R_2_ respectively:

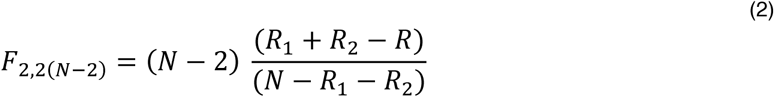

where N= N_1_ + N_2_ and R is the length of the resultant vector for the pooled direction vector observations from both groups. The resultant vector sums of all observations, R_1_, R_2_, and R, are calculated as follows:

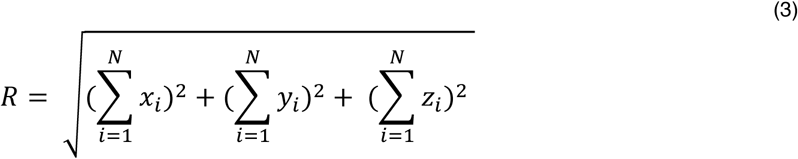

where x_i_, y_i_, z_i_ are the coordinates of the map centroids for each participant.

We performed the following F contrasts, separately for each hemisphere: controls vs patients with right CRPS and controls vs patients with left CRPS (four F tests in total). The larger the value of F, the more different the two group mean directions. A p-value was obtained using the appropriate degrees of freedom (2 and 2(N-2), respectively) and critical probability level of 0.05. The F statistics for H_0_ (no difference) and H_1_ was used to calculate the BF for each contrast, as follows (Held and Ott, 2018, equation 5):

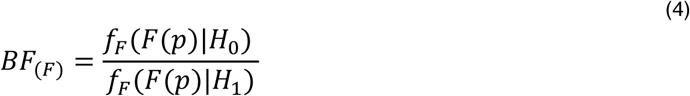

The F-based BF_10_ is simply equal to 1/BF_(F)_.

As a complementary measure of map location, we computed the geodesic distance, in mm, between the map centroid and an arbitrary reference point located within the concavity of the central sulcus (displayed in Figure 6C). Geodesic distances were statistically compared using both a frequentist and a Bayesian mixed-effects ANOVA with a within-subject factor ‘side’ (2-levels: affected, unaffected) and a between-subjects factor ‘group’ (2-levels: controls, CRPS).

Note that we did not estimate the centroid of each finger representation because our mapping method is not designed to reveal independent representations of individual fingers, given that each finger is stimulated in succession. Future studies are required to investigate finger-specific representations in CRPS.

### Evaluation of hand map geometry

As a measure of the functional geometry of the map, we measured the spatial arrangement (i.e. direction) of the spatial gradients of the map. As illustrated in Figure 7A, the hand map exhibits a typical spatial gradient from index finger to little finger. For each participant, we resampled the map ROIs from the inflated cortical surface of each participant onto a flattened, two-dimensional, surface patch. After sampling the complex-valued 3D phase-mapping data to the folded surface, we displayed it on a small flattened, 2D surface patch, which minimizes deviations from original geometry. We gently smoothed the complex values on the surface using a 1.5 mm kernel and then converted the complex-valued data (real, imaginary) to amplitude and phase angle. The 2D gradient of the phase angle was computed after fitting a plane to the data from the surrounding vertices (taking care to circularly subtract the angular data). The amplitude of the gradient at each vertex was then normalized for display.

The mean direction of map gradients is not informative because each participant cortical patch has an arbitrary direction. However, the spread (or variability) of map gradients is informative, because it doesn’t depend on the orientation of the cortical surface patch; higher variability of gradient directions indicates that the map phases are more spread and less spatially organized. Therefore, we investigated whether the functional geometry of the map is affected by CRPS, by testing whether the gradient directions of the map of the affected hand were more variable than those of the unaffected hand and controls. As a measure of map gradient variability, we calculated the circular variance of the gradient angles of each ROI. We conducted a Harrison-Kanji test (Harrison and Kanji, 1988; Berens, 2009) on the gradient variances to statistically compare the variability of map gradients across groups and participants. This test allowed us to perform a two-factor ANOVA for circular data, with a within-subject factor ‘side’ (2-levels: affected, unaffected) and a between-subjects factor ‘group’ (2-levels: controls, CRPS). BFs for each contrast were calculated as described by equation 4 (the probability level for H_0_ was 0.05).

We tested the hypothesis that there was a relation between map gradient variability and disease duration, using the equation for circular-linear correlation (r_cl_) described in (Zar, 1999: equation 27.47). A p-value for r_cl_ is computed by considering the test statistic N r_cl_, which follows a χ^2^ distribution with two degrees of freedom (Berens, 2009). BFs based on the χ^2^ distribution were calculated following equation 4 (with 0.05 probability level for H_0_).

### Cross-subject average (for illustration)

We averaged maps across subjects purely for illustration. All statistical analyses were performed on measures derived from the individual-subjects maps. We first inflated each participant’s cortical surface to a sphere, and then non-linearly morphed it into alignment with an average spherical cortical surface using FreeSurfer’s tool mri_surf2surf (Fischl et al., 1999). This procedure maximizes alignment between sulci (including the central sulcus), while minimizing metric distortions across the surface. Four steps of nearest-neighbour smoothing (<1.5 mm FWHM in 2D) were applied to the data after resampling on the spherical surface. Complex-valued mapping signals were then combined across all subjects (independently of whether the S1 map was detected or not) on a vertex-by-vertex basis by vector averaging (Mancini et al., 2012). The amplitude was normalized to 1, which prevented overrepresenting subjects with strong amplitudes. Finally, a scalar cross-subject F-ratio was calculated from the complex data and rendered back onto ‘fsaverage’ (uncorrected, p < 0.05).

### Software and Data Availability

Software to perform phase-mapping analyses is openly available at http://www.cogsci.ucsd.edu/~sereno/.tmp/dist/csurf. We used an open-source software (JASP) for the Bayesian statistical analyses: https://jasp-stats.org. Each individual hand map ROI is available at <OSF link to be disclosed upon acceptance>.

## Results

### Demographics and sensitivity to pain

Table 1 reports the demographic and clinical information of the study sample (Healthy controls: n = 17; CRPS to the left hand: n = 8; CRPS to the right hand: n = 10). Age was similar in the control group (mean ± SD, 44.9 ± 12.0 years) and in the patients (44.2 ± 11.3; independent samples t-test: t_33_ = 0.19, p = 0.856, BF_10_ = 0.329). Handedness was evaluated using the Edinburgh Handedness Inventory, which yields a laterality score that ranges from -100 (left-hand dominant) to +100 (right-hand dominant) (Oldfield, 1971). This laterality score was comparable in controls (73.6 ± 49.8) and patients (61.6 ± 58.1; independent samples t-test: t_33_ = 0.65, p = 0.518, BF_10_ = 0.384). Age of patients was similar to those found in the UK CRPS Registry (Shenker et al., 2015): mean age at onset was 43 ± 12.7 years (n = 239), whereas mean pain duration was 2.9 years (n = 237) was slightly shorter in the UK CRPS registry.

As expected, CRPS patients were more sensitive to pressure, with lower average pain pressure threshold (PPT) on their affected hand (3.4 ± 3.8) than on their unaffected hand (7.6 ± 11.0; paired samples t-test: t_17_ = -2.21, p = 0.041, BF_10_ = 1.679). Confirming that the CRPS was unilateral, PPTs on the unaffected hand of CRPS patients were similar to those of controls (average left and right hand of controls ± SD, 10.7 ± 14.9; independent samples t-test: t_33_ = 0.72, p = 0.476, BF_10_ = 0.398). Ratings of spontaneous pain did not vary in a consistent fashion before and after the imaging session (mean difference ± SD, 0.6 ± 2.5; t_16_ = 0.96, p = 0.351, BF_10_ = 0.281).

### Somatotopic representation of the hand in S1

We stimulated the tips of each finger in succession, as shown in Figure 2A, using a mechanical probe. Mechanical stimulation to the fingertips elicited a periodic response in the hand region of S1 (Figure 2B). A selection of single-subjects maps is shown in Figure 3 and the average maps are displayed in Figure 4. The map phase angle (indicating finger preference) is displayed using a continuous colour scale (red to green to blue to yellow), the saturation of which is masked by the statistical threshold. All analyses were performed on individual subject data (cluster-corrected at p < 0.01), but uncorrected group maps (p < 0.05) are displayed in Figure 4 merely for illustration. Phases corresponding to rest (no stimulation) have been truncated. The map showed a clear spatial gradient of digit preference, progressing from d2 (index finger) to d3, d4 and d5 (little finger). The arrangement and location of the map was qualitatively similar to that reported in previous human fMRI studies (Sanchez-Panchuelo et al., 2010; Mancini et al., 2012; Besle et al., 2013; Martuzzi et al., 2014; Kolasinski et al., 2016a).

**Figure 2.**
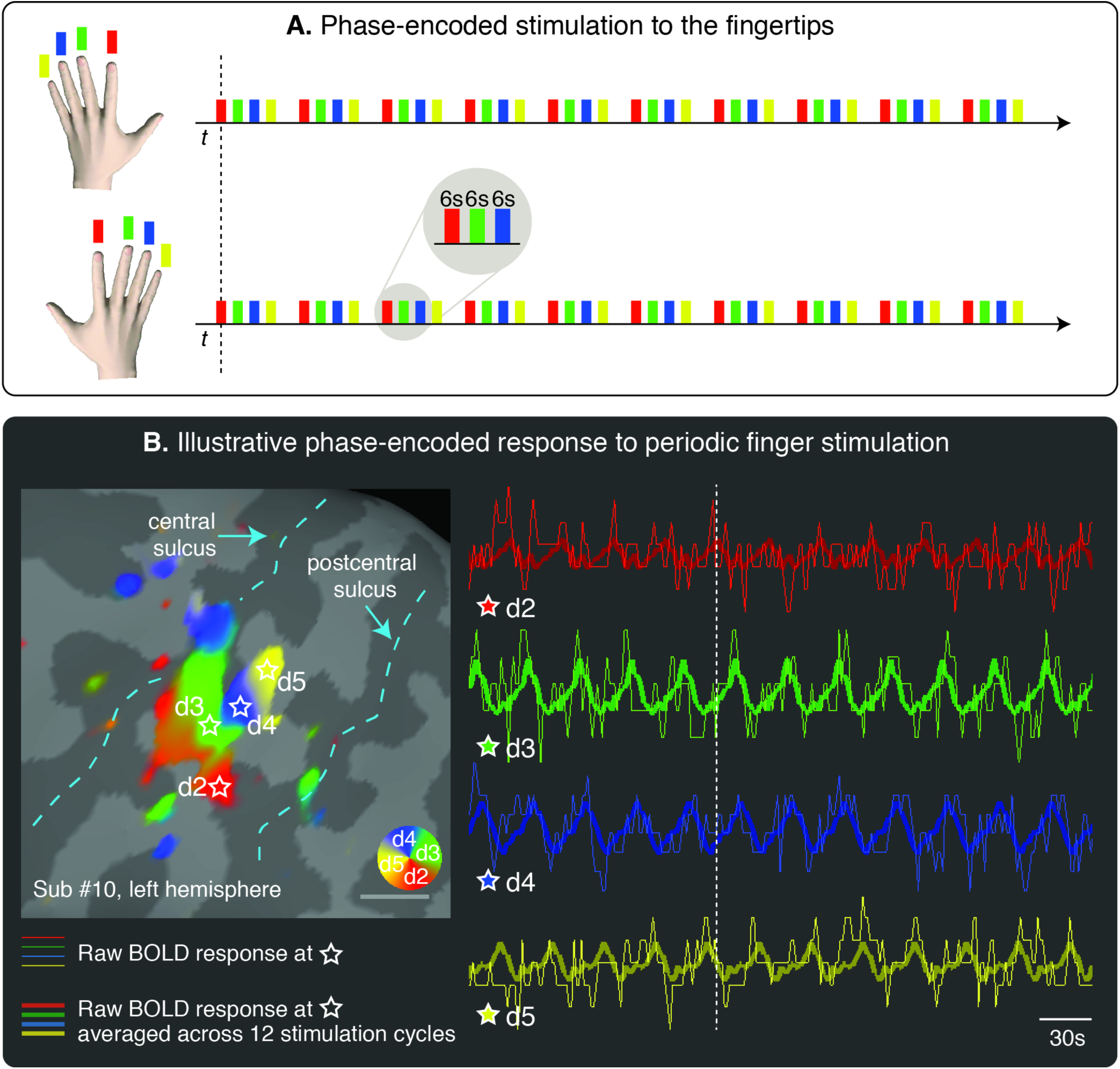
*(A) Phase-encoded stimulation procedure*. The tip of the index finger (red, d2), middle finger (green, d3), ring finger (blue, d4), little finger (yellow, d5) were stimulated in succession, in repeated cycles (12 cycles per run). To reduce scanning time, the homologous fingers of the right and left hands were stimulated simultaneously. *(B) Illustrative phase-encoded response to periodic fingertip stimulation*. The figure shows the raw Blood-Oxygen-Level-Dependent (BOLD) response in four voxels of interest (thin lines; data were motion-corrected and the linear trend removed). The locations of the voxels are marked with a star on the cortical surface of the left primary somatosensory cortex of one participant. The thicker lines represent the average of the raw BOLD response across 12 cycles of stimulation. The transversal, dashed, white line is displayed to facilitate the visualization of the shift of the phase of the BOLD response across the four voxels. The F-statistics of the signal at different phases are rendered on the inflated cortical surface and color-coded as in panel A (cluster-corrected p < 0.01). Phases corresponding to rest have been truncated.

**Figure 3.**
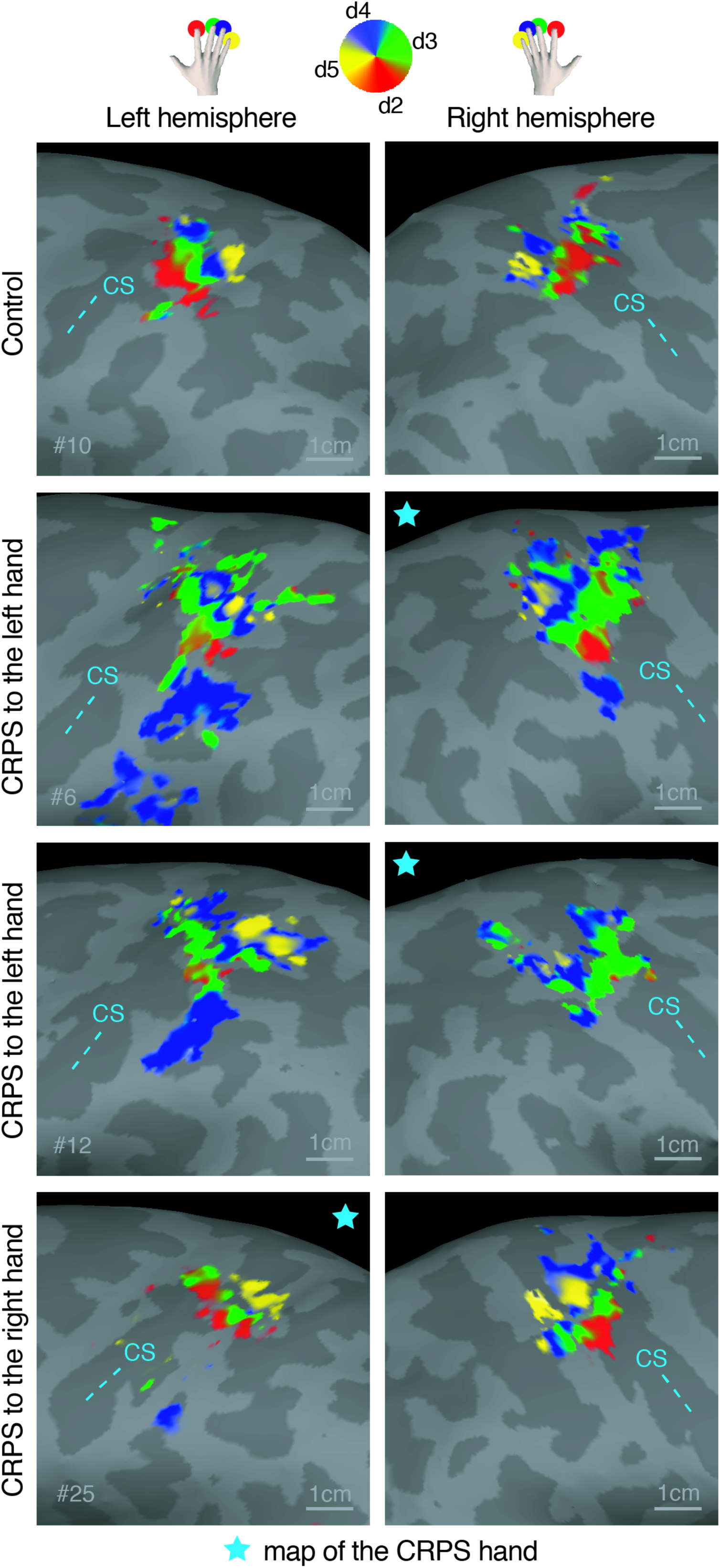
*Phase maps of the hand in an illustrative control participant and three CRPS patients.* The color-coding scheme used is shown on the top of the figure and is the same as in Figure 1: red = d2, green = d3, blue = d4, yellow = d5. Phases corresponding to rest have been truncated. Statistical thresholding and cluster correction at p < 0.01 was applied to each individual-participant data. CS: central sulcus. The star symbol denotates the map of the CRPS hand.

**Figure 4.**
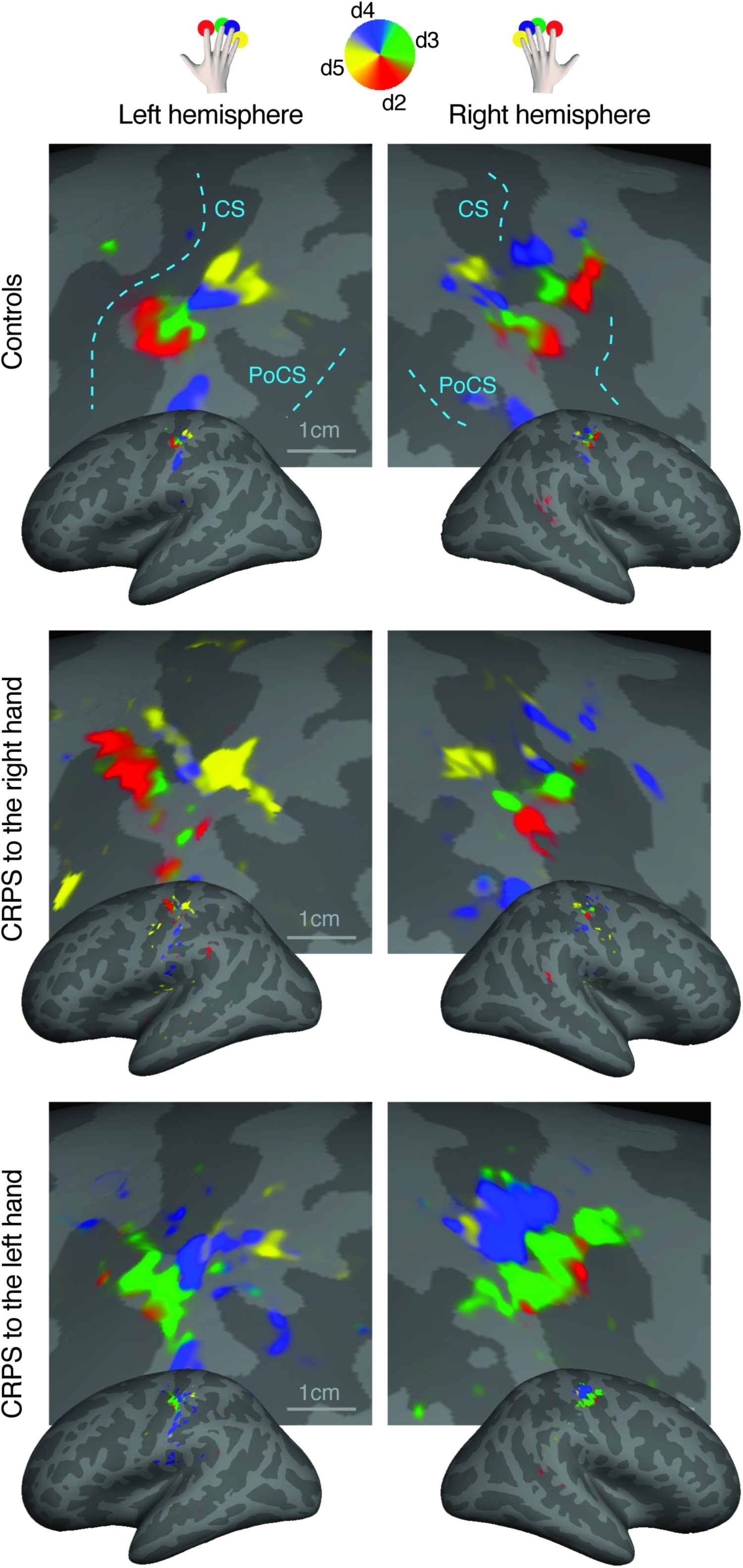
*Surface-based average of phase maps in controls, patients with CRPS to the right hand, and patients with CRPS to the left hand.* The complex-valued mapping data were averaged in a spherical surface coordinate system after morphing each subject’s data into alignment with an average spherical sulcal pattern, and the F-statistics were rendered back onto an average unfolded cortical surface (Freesurfer’s fsaverage, ‘inflated_average’; uncorrected p < 0.05 only for illustration). The color-coding scheme used is shown on the top of the figure and is the same as in Figures 1-2: red = d2, green = d3, blue = d4, yellow = d5. Phases corresponding to rest have been truncated. CS: central sulcus; PoCS: post-central sulcus.

We tested whether the area, location, and functional geometry of the map of the affected hand was similar to those of the unaffected hand and controls. To do so, we defined individual ROIs as clusters located in S1 that showed a significant periodic response at the spatial frequency of stimulation (cluster-corrected, p < 0.01).

#### 1. Map area

To control for inter-individual variability in brain size, we resampled the phase maps onto the original average brain volume. We then calculated the surface area of the left- and right-hand maps from each participant ROI. In order to increase statistical power, we flipped the data from the right hand CRPS group so that the affected side became the left hand/right hemisphere in all patients and then pooled these data. As evident in Figure 5A, the map area was comparable among groups and sides. A mixed-effects ANOVA with a within-subject factor ‘side’ (2-levels: affected, unaffected) and a between-subjects factor ‘group’ (2-levels: controls, CRPS) did not provide evidence for any main effect or interaction (‘side’: F_1,28_ = 0.281, *p* = 0.60, _p_η^2^ = 0.010; ‘group’: F_1,28_ = 1.555, *p* = 0.223, _p_η^2^ = 0.053; ‘side’ by ‘group’: F_1,28_ = 0.315, *p* = 0.579, _p_η^2^ = 0.011). A Bayesian mixed-effects ANOVA provided the stronger evidence for the null model (BF_10_ = 1, P(M|data) = 0.492) relative to models of ‘group’ (BF_10_ = 0.512, P(M|data) = 0.252), ‘side’ (BF_10_ = 0.305, P(M|data) = 0.150), ‘side+group’ (BF_10_ = 0.153, P(M|data) = 0.075), ‘side+group+interaction’ (BF_10_ = 0.126, P(M|data) = 0.030).

**Figure 5.**
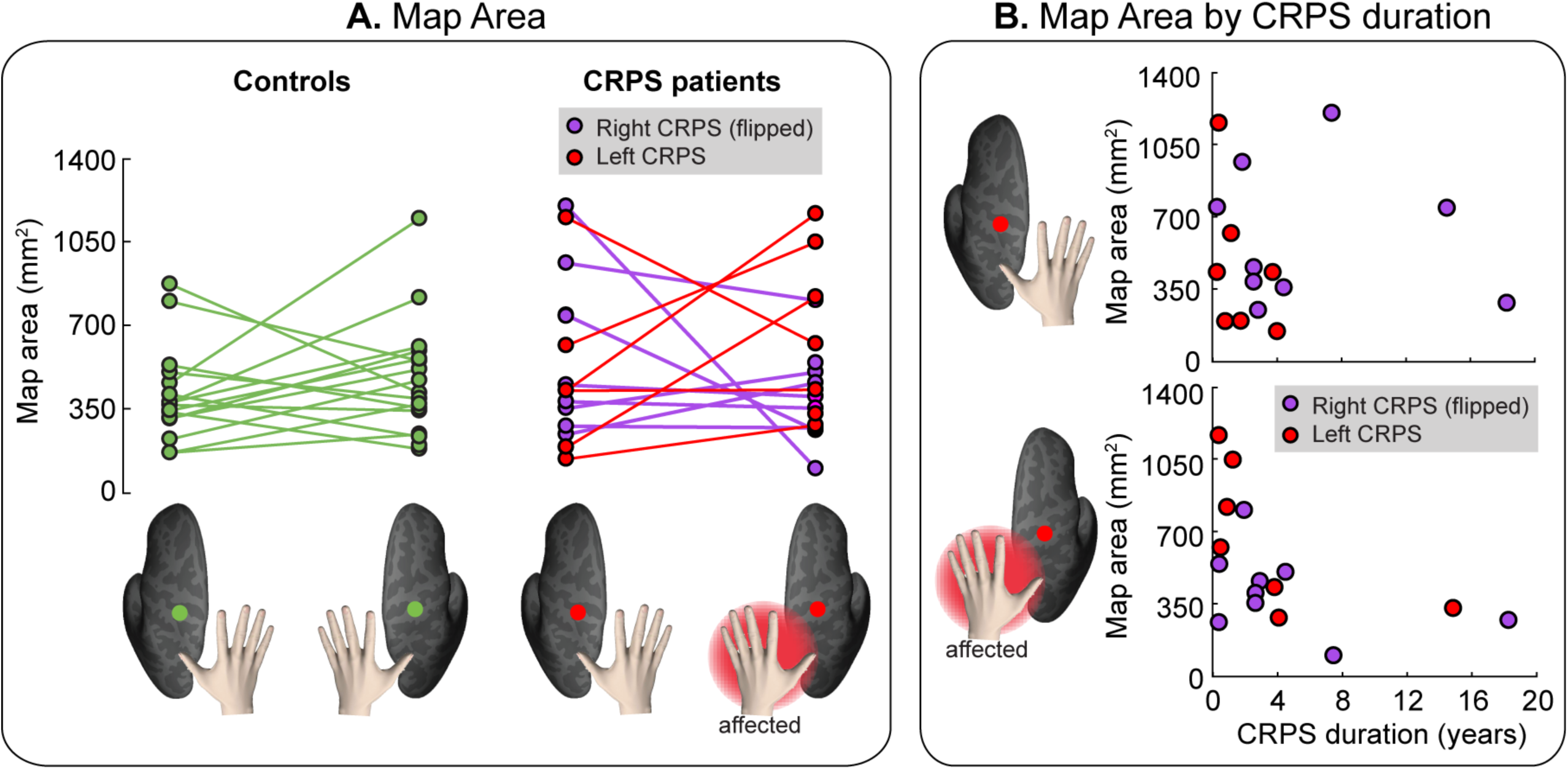
*(A) Area of the hand map in S1*. The area of the hand map (mm^2^) in the left hemisphere and right hemisphere is plotted for each group and individual participant. To facilitate comparison, data from the two CRPS groups (right hand CRPS, left hand CRPS) were pooled, after flipping the data from one group (right hand CRPS) so that the affected side is the left hand/right hemisphere in all patients. *(B) No relation between map area and CRPS duration*. The top plot shows the lack of relation between the S1 map of the healthy hand and CRPS duration, whereas the bottom plot shows the weak (not significant) relation between the S1 map of the affected hand and disease duration.

**Figure 6.**
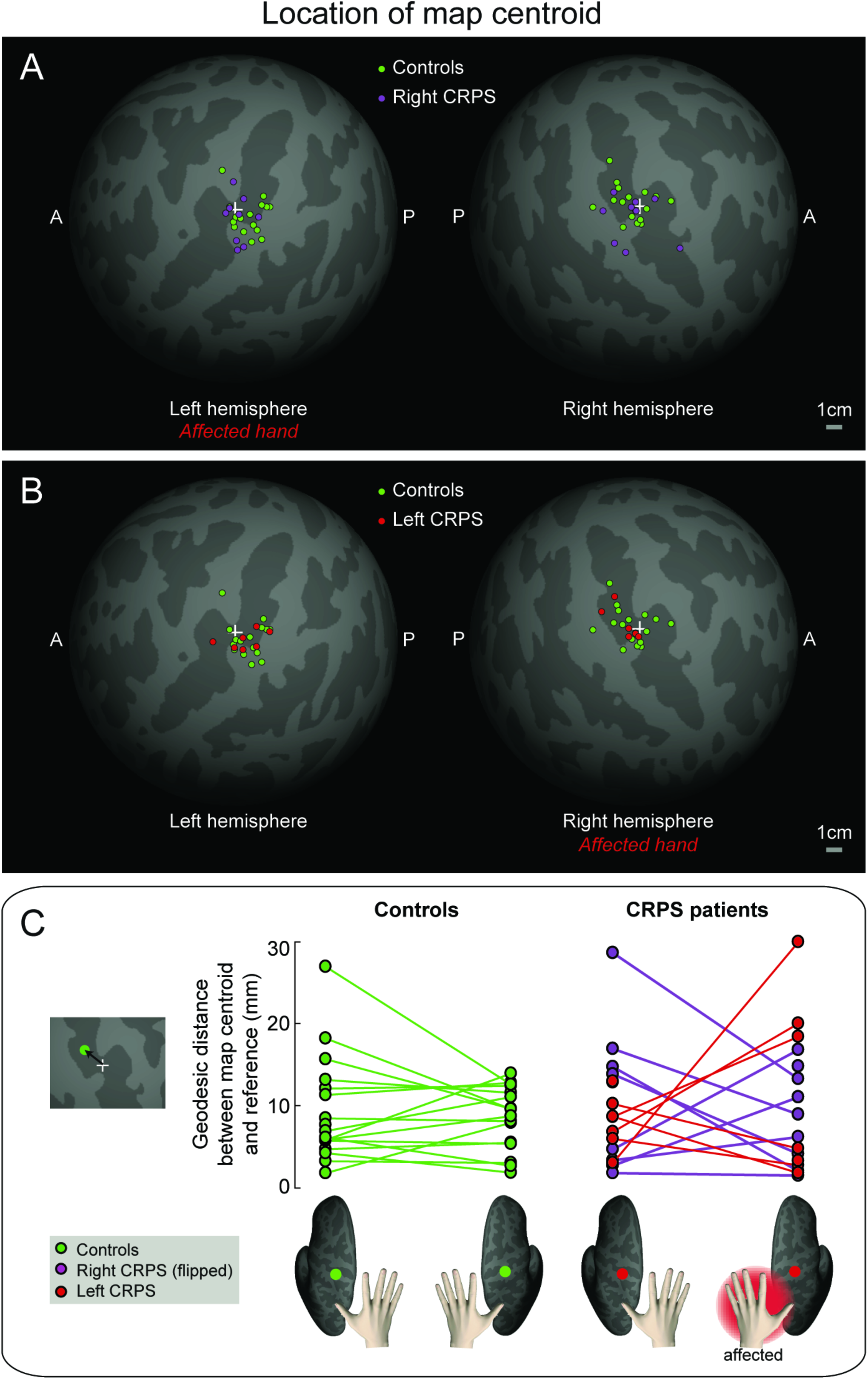
*(A-B) Spatial distribution of map centroids.* The location of the centroid of the hand map in each individual subject is displayed on an average spherical cortical surface. An arbitrary reference point on the central sulcus is marked with a white cross. *(C) Geodesic distance (mm) between each map centroid and a reference point (‘+’) on the central sulcus*. To facilitate comparison, data from the two CRPS groups (right hand CRPS, left hand CRPS) were pooled, after flipping the data from one group (right hand CRPS) so that the affected side is the left hand/right hemisphere in all patients.

**Figure 7.**
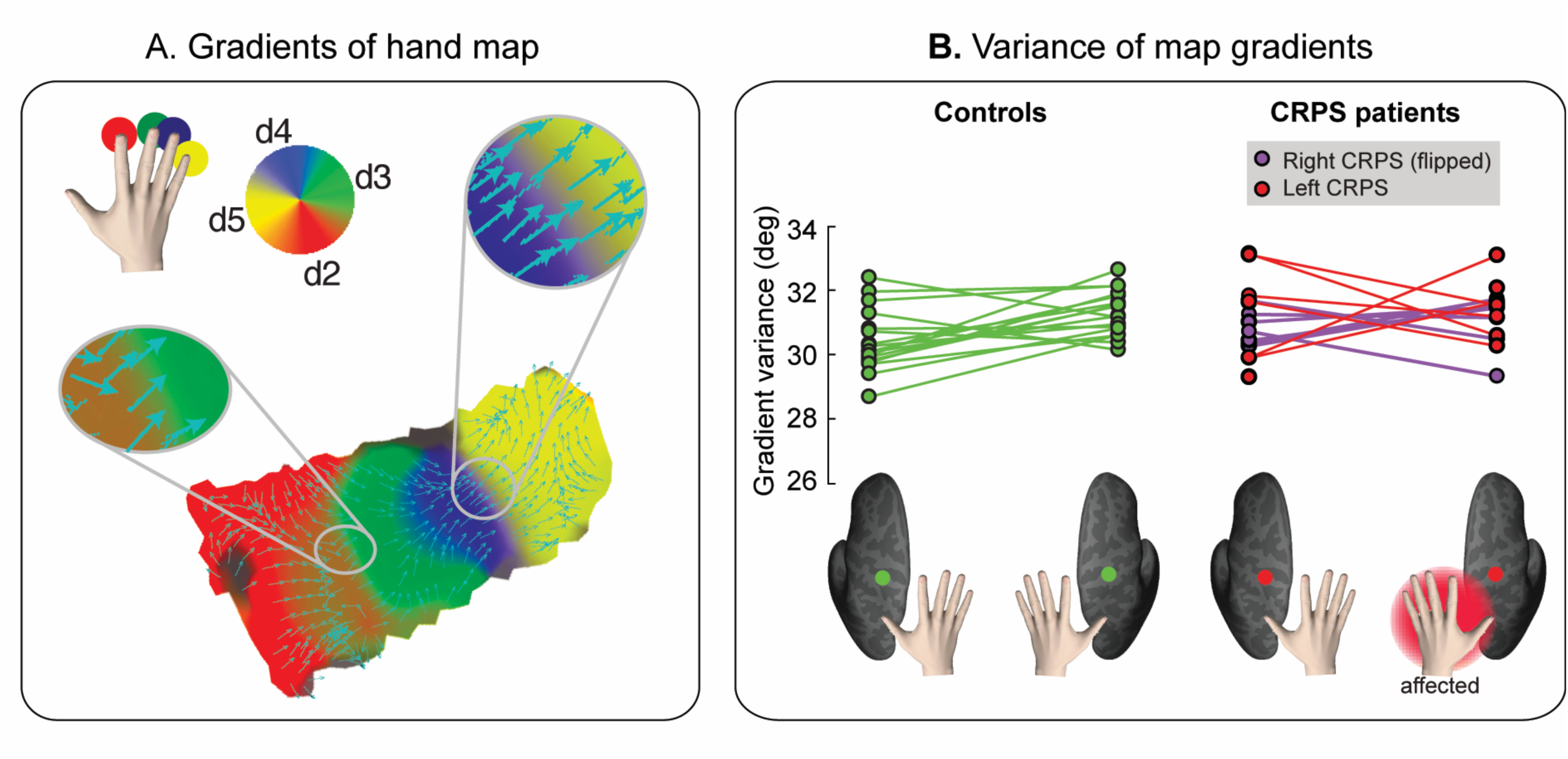
*(A) Gradients of the hand map*. Gradients of a single-subject phase map are displayed as cyan arrows over a flattened (2D) cortical surface patch. The gradient points in the direction of the greatest rate of increase of the function (i.e. the direction of the phase shift in the hand map). The color-coding scheme of the hand map is the same as in Figures 1-3: red = d2, green = d3, blue = d4, yellow = d5. *(B) Variability of hand map gradients*. The circular variance of map gradient directions is displayed for each participant and condition (side: left hemisphere, right hemisphere; group: controls, CRPS patients). The color-coding scheme for panel B is shown at the bottom of the figure. To facilitate comparison, data from the two CRPS groups (right hand CRPS, left hand CRPS) were pooled, after flipping the data from one group (right hand CRPS) so that the affected side is the left hand/right hemisphere in all patients.

#### 1.1. Relation with disease duration

We checked whether there was any relationship between map size and duration of disease. As the distribution of disease duration values was skewed towards small values, disease duration data were first transformed onto natural logarithms. The log-transformed disease duration was normally distributed (Shapiro-Wilk test = 0.94, p = 0.293; skewness = -0.002 ± 0.536). A Bayesian linear regression showed stronger evidence for a model in which the area of the map covaried with (log-transformed) disease duration BF_10_ = 3.016, P(M|data) = 0.030) than for the null model (BF_10_ = 1, P(M|data) = 0.249): in participants that suffered from CRPS for a longer time had a smaller map of the affected hand.

#### 1.2. Relation with upper limb disability

Disease duration did not correlate with the severity of disability of the CRPS limb, as measured by the *Quick*Dash score (Pearson’s r = 0.002, BF_10_ = 0.291). In patients, the QuickDash score did not predict the area of the map of the affected hand in a Bayesian linear regression analysis (null model: BF_10_ = 1, P(M|data) = 0.620; disability model: BF_10_ = 0.614, P(M|data) = 0.380).

#### 1.3. Relation with pain intensity

We found no evidence for a linear relation between the area of the map of the affected hand and pain intensity ratings obtained during the imaging session (ratings for each subject are reported in Table 1). The null model (BF_10_ = 1, P(M|data) = 0.685) won over a model in which the area of the map of the affected hand linearly covaried with pain intensity ratings (BF_10_ = 0.460, P(M|data) = 0.315). Moreover, there was no evidence for a relation between the area of the map of the affected hand and a score of pain severity derived from PPTs (PP_severity_). A Bayesian linear regression showed that the null model (BF_10_ = 1, P(M|data) = 0.510) and a model with PP_severity_ (BF_10_ = 0.960, P(M|data) = 0.490) were similarly likely. In our sample, disease duration did not correlate with either pain intensity ratings collected during the imaging session (Pearson’s r = 0.249, BF_10_ = 0.463) or with PP_severity_ (Pearson’s r = -0.108, BF_10_ = 0.317)

Finally, the area of the map of the unaffected hand was not explained by CRPS duration, pain intensity rating, or PP_severity_ (all BF_10_ for null models = 1; all BF_10_ for alternative models < 0.6).

In summary, these analyses do not provide support for the hypothesis that the map of the CRPS hand was smaller than the map of the unaffected hand and that of healthy controls, at group level. However, we found evidence that the map area of the CRPS hand was modulated by disease duration, across participants. The more chronic was the disease, the smaller was the map of the affected hand. Map area was not predicted by various measures of pain severity and upper limb disability.

#### 2. Map location

We calculated the centroid of the hand map, after resampling it onto an average spherical surface (see “Evaluation of hand map location” for details). This was done to control for individual differences in brain morphology and to obtain localisation measures that were not confounded by gyrification. Figure 6A-B shows the distribution of map centroids of each participant, resampled onto a canonical spherical cortical surface of an average brain; the map centroid location was variable among participants of each group, but visibly similar across groups. Indeed, the F-statistics based on the Fisher probability density function (Fisher, 1953) did not provide evidence for any directional difference between groups for either side (Table 2).

**Table 2.** Statistical values for the comparison of the locations of map centroids between groups.

| **Contrast** | **Side** | **F** | **Degrees of freedom** | **p** | **BF<sub>10</sub>** |
| --- | --- | --- | --- | --- | --- |
| Controls vs Right CRPS | Left hemisphere | 0.002 | 2,50 | 0.998 | 0.001 |
| Controls vs Right CRPS | Right hemisphere | 0.025 | 2,48 | 0.975 | 0.306 |
| Controls vs Left CRPS | Left hemisphere | 0.005 | 2,42 | 0.995 | 0.309 |
| Controls vs Left CRPS | Right hemisphere | 0.001 | 2,44 | 0.999 | 0.311 |

As a further comparison of the locations of map centroids across groups, we computed the geodesic distance, in mm, between the map centroid and an arbitrary reference point located within the concavity of the central sulcus (Figure 6C). Importantly, geodesic distance measures calculated onto average spherical surfaces are not confounded by gyrification and allow comparison of different subjects. This is a key advantage of our approach over previous studies which measured Euclidean distances between two finger representations. A mixed-effects ANOVA with a within-subject factor ‘side’ and a between-subjects factor ‘group’ did not provide evidence for any main effect or interaction (‘side’: F_1,28_ < 0.01, *p* = 0.999, _p_η^2^ < 0.001; ‘group’: F_1,28_ = 0.163, *p* = 0.689, _p_η^2^ = 0.006; ‘side’ by ‘group’: F_1,28_ = 0.254, *p* = 0.619, _p_η^2^ = 0.009). In a Bayesian mixed-effects ANOVA, the null model had stronger evidence (BF_10_ = 1, P(M|data) = 0.581) relative to models of ‘group’ (BF_10_ = 0.340, P(M|data) = 0.198), ‘side’ (BF_10_ = 0.262, P(M|data) = 0.152), ‘side+group’ (BF_10_ = 0.087, P(M|data) = 0.051), ‘side+group+interaction’ (BF_10_ = 0.032, P(M|data) = 0.019).

Altogether, these analyses indicate that the location of the hand map centroid was not affected by CRPS.

#### 3. Map geometry

Finally, we evaluated the variability of the geometry of the map of the affected hand in CRPS patients. As illustrated in Figure 7A, the hand map exhibits a typical spatial gradient from index finger to little finger. The spatial gradient (i.e. the direction) of the map indicates the spatial progression of the map phases, providing a measure of the map geometry. We investigated whether the gradient directions of the map of the affected hand were more variable than those of the unaffected hand and controls. As a measure of map gradient variability, we calculated the circular variance of the gradient angles of each flattened, two-dimensional, surface ROI (see “Evaluation of hand map geometry” for details).

The gradient directions of the map of the affected hand were not differently variable (i.e. not differently spread) from those of the unaffected hand and controls (Figure 7B). A Harrison-Kanji test with a within-subject factor ‘side’ and a between-subjects factor ‘group’ on the gradient variances provided evidence for a main effect of side (F_1,59_ = 4.813, *p* = 0.032, _p_η^2^ = 0.079, BF_10_ = 1.202) and no evidence for a main effect of group (F_1,59_ = 2.243, *p* = 0.140, _p_η^2^ = 0.038, BF_10_ = 0.560). We found weak and inconclusive evidence for an interaction between side and group (F_1,59_ = 3.889, *p* = 0.071, _p_η^2^ = 0.057, BF_10_ = 0.971). This suggests that the spread of map gradients, which is a measure of functional organization, was largely similar across groups.

Lastly, we tested whether there was a circular-linear correlation (r_cl_) between map gradient variability and disease duration. This analysis did not provide evidence for a relation between the (log-transformed) disease duration and the gradient variability of either the affected hand map (r_cl_ = 0.395, *p* = 0.310, BF_10_ = 0.066) or the unaffected hand map (r_cl_ = 0.197, *p* = 0.734, BF_10_ = 0.034).

## Discussion

We show that the cortical map of the fingertips of the CRPS hand in S1 is strikingly comparable to the map of the unaffected hand and controls in terms of area, location, orientation, and geometry. Although the area of the map of the affected hand was comparable to that of the unaffected hand and controls, it was modulated by disease duration, but not by pain intensity, pain sensitivity and upper limb disability: across participants, the longer was the duration of CRPS, the smaller was the area of the map of the CRPS hand. Our results do not exclude that other abnormalities may occur at S1 level, such as excitability changes (Lenz et al., 2011; Di Pietro et al., 2013), morphological (Baliki et al., 2011; Pleger et al., 2014; cfr. van Velzen et al., 2016) and connectivity changes (Geha et al., 2008). However, our findings challenge or, at the very least, narrow the notion of S1 map reorganization in CRPS: if any map reorganization occurs, it does not appear to be directly related to pain.

These findings urge us to reconsider the mechanisms that are proposed to underpin CRPS (Marinus et al., 2011). They also compel us to revaluate the rationale for (and mechanism of effect of) clinical interventions that aimed to reduce pain by “restoring” somatotopic representations with sensory discrimination training (Moseley et al., 2008b; Catley et al., 2014), or by correcting sensorimotor incongruences (which are thought to be induced by S1 reorganisation) with mirror therapy (McCabe et al., 2003) (but see Moseley and Gandevia, 2005; Moseley et al., 2008b). Although these interventions appear to offer clinical benefit (O’Connell et al., 2013), they are unlikely to engender a “restoration” of somatotopic representations in S1, which are largely comparable to those of controls.

### Revisiting previous evidence of somatotopic reorganisation in CRPS

Comparisons across different studies are inevitably challenging due to the complexity and variety of CRPS symptomatology; in previous studies, patients varied greatly in regard to the combination, severity and duration of their symptoms. Our study suggests that map size is probably related with disease duration, although only a longitudinal study could confirm a causal relationship.

There are also important methodological issues to consider. The notion of somatotopic reorganisation in CRPS was mostly based on studies that used imaging methods (EEG/MEG) with lower spatial resolution than fMRI (Juottonen et al., 2002; Maihofner et al., 2003; Pleger et al., 2004; Vartiainen et al., 2008; Vartiainen et al., 2009). A more recent study used fMRI and measured the cortical distance between d1 and d5 activation peaks (Di Pietro et al., 2015). This study partially confirmed former EEG/MEG findings, reporting that the d1-d5 distance in S1 was smaller for the affected hand than it was for the unaffected hand in CRPS patients. However, the representation of the affected hand was comparable to that of healthy controls, in agreement with the current results. Critically, the Di Pietro study (2015) found that the representation of the unaffected hand in CRPS patients was larger than that of controls, thus challenging the view that the representation of the affected hand is shrunk and suggesting that the representation of the unaffected hand is actually enlarged. The current results do not support either interpretation.

Three important limitations affect all previous studies, regardless of the imaging approach used. First, the approach taken to estimate map size is both indirect and incomplete, because it is based on the measurement of the Euclidean *distance* between the activation maxima of two fingers (d1 and d5). Instead, the *area* of the map of all fingers is a more direct and complete measure of map size. Second, Euclidean measures of cortical distances can be inaccurate because they disregard that the cortical surface is not flat, especially in the regions of the sulci. Third, Euclidean distance measures can be affected by non-topographical, structural changes in SI, which can be associated with CRPS (Baliki et al., 2011; Pleger et al., 2014). The latter two problems can be overcome by morphing activation maps onto a reconstruction of the flattened cortical surface (Makin et al., 2013a; Kikkert et al., 2016), but previous studies on CRPS patients have not taken this approach. Altogether, these methodological issues can affect both the accuracy and validity of previous measures of map extent.

### Stability of cortical topographies

Recent fMRI studies (Makin et al., 2013a; Kikkert et al., 2016) suggest that finger topographies in S1 are surprisingly persistent even in humans who suffered amputation of the upper-limb. It was demonstrated that the area, location and functional organisation of the S1 maps of the missing hand were similar, although noisier, to those observed in controls during finger movements (Makin et al., 2013a; Kikkert et al., 2016). It has also been shown that the deafferented territory in human S1 can respond to somatotopically adjacent body regions (i.e. the lip for upper limb amputees) (Flor et al., 1995; Flor, 2008), or to body regions that the amputees overuse to supplement lost hand function (e.g. the intact hand). This results in a highly idiosyncratic remapping which does not necessarily involve adjacent representations in S1 (Makin et al., 2013b; Philip and Frey, 2014). Thus, cortical reorganisation in amputees is not dictated by cortical topographies, but can depend on compensatory use of other body parts. Similarly, short-term shifts in S1 maps can occur in healthy participants after surgical gluing of the index and middle fingers for 24 hours. These changes are thought to depend on compensatory use of the fourth and fifth fingers (Kolasinski et al., 2016b). These studies support the view that any S1 change previously reported in CRPS patients might not directly related to pain, but it remains to be determined why map shrinkage relates to disease duration. Could it be related to hand use? We found no relation between map size and severity of the upper limb disability.

Recent evidence from electrophysiological and inactivation studies in monkeys suggests that the reorganisation following nerve transection originates, not in S1, but in the brainstem. Indeed, inactivating the cuneate nucleus abolishes the neural activity in the deafferented limb representation in S1 elicited by face stimulation (Kambi et al., 2014). Hence, loss of input from a body region in adulthood may lead to the formation or potentiation of lateral connections in the brainstem, which gives rise to a new pathway from periphery to cortex. It is not clear whether this new pathway contributes to cortical reorganisation, but the original pathway seems to be relatively spared even under the extreme circumstance of limb amputation (Makin and Bensmaia, 2017).

Some resistance to change has also been described for visual retinotopic maps. Although it has been shown that large lesions to the retina in adult mammals can induce a reorganization of retinotopic cortical maps in primary visual cortex (Kaas et al., 1990), more recent studies have reported that the topography of the macaque primary visual cortex does not change (for at least seven months) following binocular retinal lesions (Smirnakis et al., 2005). Similarly, severe eye diseases such as retinal degeneration do not seem to affect retinotopic representations in the human early visual cortex (Xie et al., 2012; Haak et al., 2016). Altogether, these findings suggest that cortical topography is more stable and resistant to change than what it was initially thought.

### Conclusion and future directions

Our study provides the most complete characterization, to date, of the S1 somatotopy of the CRPS hand. We report that the S1 representation of the CRPS hand is comparable, at the group level, to that of the healthy hand, in terms of cortical area, location and geometry. The area of the S1 map of the CRPS hand is related to disease duration but not to pain intensity, pain sensitivity and upper limb disability. Future longitudinal studies are required to determine how the map changes over time and its effect on sensorimotor function.

## Author contributions

FM, APW, MMS, JHM, GDI, MIS, GLM, CR conceived and designed the study. APW, MMS, ZJI collected the neuroimaging data. APW analysed the clinical data. FM analysed the imaging data. FM wrote the initial draft of the manuscript. All authors edited the manuscript.

### Acknowledgments

We acknowledge the support of the Australian National Imaging Facility, a National Collaborative Research Infrastructure Strategy (NCRIS) capability. We are grateful to Dr Michael Green and to the staff of NeuRA Imaging. FM and GDI were supported by a Wellcome Trust Strategic Award (COLL JLARAXR). GDI was additionally supported by a ERC Consolidator Grant (PAINSTRAT). APW was supported by educational grants from the Australian Pain Society/Australian Pain Relief Association, Mundipharma (PhD Scholar #3) and NeuRA. GLM was supported by a research fellowship from the National Health and Medical Research Council (NHMRC) of Australia (ID 1061279). This study was supported by a project grant from the NHMRC (ID 630431).

